# Model-based aversive learning in humans is supported by preferential task state reactivation

**DOI:** 10.1101/2020.11.30.404491

**Authors:** Toby Wise, Yunzhe Liu, Fatima Chowdhury, Raymond J. Dolan

**Affiliations:** Max Planck UCL Centre for Computational Psychiatry and Ageing Research, University College London, London, UK; Wellcome Centre for Human Neuroimaging, University College London, London, UK; Division of the Humanities and Social Sciences, California Institute of Technology, Pasadena, CA, USA

**Author notes:** Contact: Toby Wise. These authors contributed equally (Toby Wise, Yunzhe Liu).

## Abstract

Harm avoidance is critical for survival, yet little is known regarding the underlying neural mechanisms supporting avoidance when we cannot rely on direct trial and error experience. Neural reactivation, and sequential replay, have emerged as potential candidate mechanisms. Here, during an aversive learning task, in conjunction with magnetoencephalography, we show prospective and retrospective reactivation for planning and learning respectively, coupled to evidence for sequential replay. Specifically, when subjects plan in an aversive context, we find preferential reactivation of subsequently chosen goal states and sequential replay of the preceding path. This reactivation was associated with greater hippocampal theta power. At outcome receipt, unchosen goal states are reactivated regardless of outcome valence. However, replay of paths leading to goal states was directionally modulated by outcome valence, with aversive outcomes leading to stronger reverse replay compared to safe outcomes. Our findings suggest that avoidance behaviour involves simulation of alternative future and past outcome states through hippocampally-mediated reactivation and replay.

## Introduction

Simulation of future states has gained increasing attention as a mechanism supporting decision making, particularly in the absence of direct experience^1^, This process is thought to be supported by neural replay, evident in observations that hippocampal place cells and adjacent place fields reactivate in a forward or reverse sequence, reflecting future or past trajectories^2,3^. Notably, paths leading to aversive outcomes show preferential replay during avoidance behaviour^4^, implicating a simulation of negative outcomes as a neural signature of harm avoidance. Intriguingly, one recent proposal suggests the brain’s ability to simulate both experienced and hypothetical outcomes might provide for a mechanistic understanding of phenomena such as worry and rumination^5–7^.

In humans, reactivation of individual rewarded states has been found to support planning and inference^8–11^ as well as updating of reward values^12^. Recent human neuroimaging findings show that, across a variety of tasks, reactivation of singular events can occur in sequence in a manner reminiscent of rodent hippocampal replay^13–18^. Collectively, these studies suggest non-local reactivation is a motif of human model-based learning and decision making. While a recent study demonstrated reactivation of potential outcomes during avoidance^19^, it is unclear as to what extent this is related to non-local, model-based avoidance, which is critical for successful avoidance of danger and arguably has the clearest relevance to common symptoms of mood disorders^5,20–22^.

Here our focus is on the contribution of reactivation and replay (i.e., sequential reactivation) to human avoidance behaviour. We used magnetoencephalography (MEG) to index states while subjects engaged in an aversive learning task that promotes avoidance decisions based on inference about learned aversive state values. Specifically, we ask whether non-local reactivation and replay of task states underpin model-based avoidance and aversive value updating. Our results indicate that replay supports model-based aversive learning through simulation of states that have not been the object of direct experience. A simulation of aversive states offers the intriguing possibility that this type of process might be relevant for ruminative negative thought patterns that are seen in mood disorders^7^.

## Results

### Subjects adaptively use model-based control to facilitate avoidance

28 subjects (20 female, 8 male) completed an aversive learning task while we acquired simultaneous neural data using MEG. The task space consisted 14 discrete states, each represented by unique visual images. Subjects navigated from start to terminal states (Figure 1A) associated with a drifting probability of an electric shock. Shock probabilities were designed to be moderately, but not perfectly, anticorrelated (*r*=-.57; Figure 1C). This ensured that one option was generally preferable while requiring representation of that outcome types (shock and safety) for each terminal state. Importantly, the task space included two arms (referred to as generalisation arms) that terminated at the very same state as one of the other two arms (referred to as learning arms).

**Figure 1.**
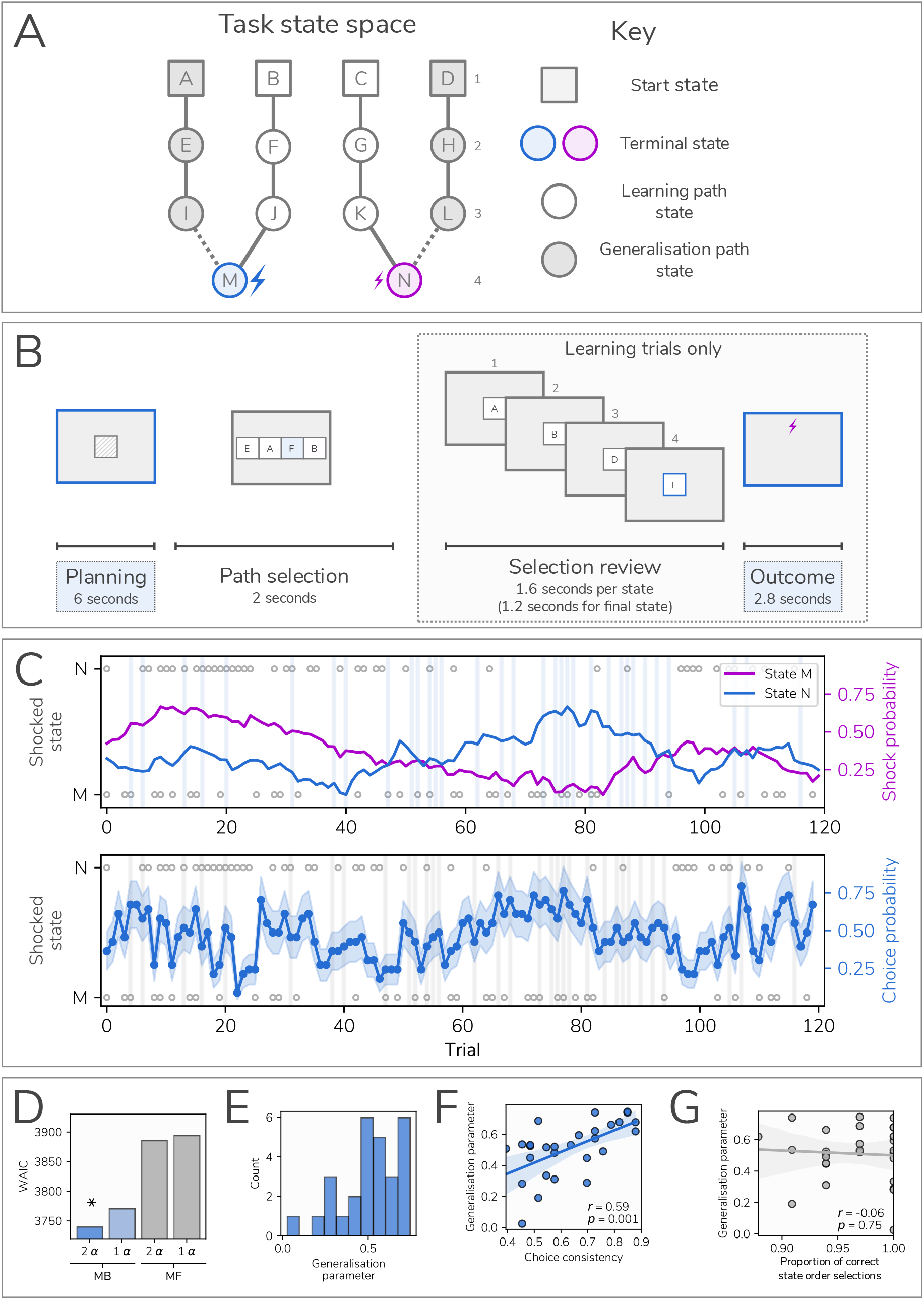
Task design and behaviour. A) Illustration of task states and transitions. Subjects navigated a map comprising 14 states (labelled A-N), each represented by a unique visual image. On “learning” trials, subjects chose between two “learning” paths (B>F>J>M, C>G>K>N), that at the terminal state led to a shock or safe outcome according to a drifting shock probability determined by a random walk. On “generalisation” trials (28%), subjects chose between the two “generalisation” paths (A>E>I>M, D>H>L>N). For choices on these paths the associated outcomes were not shown to subjects. B) Trial procedure. Each trial began with a 6 second “planning” phase during which subjects viewed a coloured square representing the trial type (one colour indicating a learning trial and another indicating a generalisation trial). Subjects were instructed to think about the sequence of states they would like to select. Subjects then selected a sequence of states that took them from a starting state (e.g. A) to one of the final states (the “state selection” phase). Subjects were presented with a state from an array of four presented images which included valid state(s) (i.e. states to which subjects could validly transition from the current selected state) as well as randomly selected invalid states. On learning trials, after selecting a path the entire sequence trajectory was shown sequentially (the “selection review”). At the final state, subjects saw either a shock icon (indicating an upcoming shock) or a crossed shock icon (indicating safety). Outcomes (both shock and safety) were accumulated and three of these were randomly administered at the end of each block of 20 trials. On generalisation trials, the trial ended after state selection without playback of the path, with subjects told the hidden outcomes would accumulate for administration upon completion of the entire task, unlike those from learning trials which were administered at the end of each block. C) Trial outcomes and subjects’ responses. In the top panel, the purple and blue lines indicate the shock probability for the final states of each learning arm; these followed independent (moderately anticorrelated) random walks, created such that each state was safest on an approximately equal number of trials. Blue vertical bars represent generalisation trials. In the bottom panel, the blue line represents the proportion of subjects choosing option N, with shaded area representing the standard error of this proportion. Grey circles indicate which state was shocked on a given trial (assuming a subject chose this state), with those at the top indicating a shock for state N and the bottom indicating a shock for state M. D) Model comparison (see Methods for description of the models) demonstrated superior performance of a model incorporating asymmetric updating from shock and safety outcomes, as well as model-based inference on generalisation trials. MF represents model-free control; MB represents model-based inference on generalisation trials. α refers to the learning rate parameter, either dependent on outcome valence (2 α) or the same for both shock and no shock outcomes (1 α). E) Estimated model generalisation parameter values across subjects. Values of 0 represent no model-based inference (i.e. choosing randomly on generalisation trials) while a value of 1 indicates choices fully consistent with that expected if these were made based on a learned value. F) Correlation between generalisation parameter values and choice consistency between adjacent learning and generalisation trials, a model-agnostic approximation of a subject’s tendency to use model-based inference. The strong relationship indicates this parameter provides a valid representation of this behaviour. G) Non-significant correlation between generalisation value parameter and the number of errors made on generalisation trials across subjects (where subjects failed to enter a correct sequence of states), showing that low generalisation parameter values do not reflect poor knowledge of task structure. If this were the case, more errors would be associated with less generalisation, which is not seen in the data.

On most trials, subjects had to choose between the two learning arms and observed an outcome screen upon reaching the terminal state. Rather than administering shocks during actual trials the outcome types accumulated over the course of each block. Subsequently, 3 randomly selected outcomes were administered at the end of a block, a design feature that avoids contamination of task phases by shock-related signals. Crucially, on a subset of trials (28% of trials, referred to as generalisation trials) subjects chose between the two generalisation paths. The outcome for a generalisation path was never shown, obviating learning through direct experience within these arms. Instead, the next trial started immediately. Subjects received prior instruction that the hidden outcomes from these generalisation trials would nevertheless accumulate and be delivered at the end of the task, ensuring decisions on generalisation arms promoted model-based inference in relation to an hypothetical final state.

To examine the computational processes underlying task avoidance behaviour, we fit a reinforcement learning model that updates aversive value on each trial according to an experienced prediction error^23^. Learning was weighted via separate learning rates for better and worse than expected outcomes, allowing an asymmetry in learning previously observed in these types of aversive learning tasks^24,25^. This provided a better fit to the data, as assessed using the Watanabe-Akaike Information Criterion (WAIC)^26^, than a variant model relying on symmetric updating (asymmetric WAIC = 3744.55, symmetric WAIC = 3770.09, Figure 1D). As in previous work, subjects varied in their tendency to learn from safety versus punishment^27^ but we found no consistent bias towards learning from danger when comparing learning rates for shock and safety outcomes (t(27) = 0.85, *p* = 0.40), as observed in our prior studies^24,25^.

Importantly, we incorporated into our model an additional mechanism that allowed learned values for the two end states to inform decisions on generalisation trials, which rely solely on the exercise of model-based inference (see methods). The degree of model-based inference was represented by a free parameter *G*, where a value 0 entailed choices on generalisation trials were random and a value of 1 entailed choice probability was identical to that for learning trials. Model comparison indicated that models including model-based inference outperformed models that chose randomly on generalisation trials (WAIC for best performing model without generalisation = 3888.18), consistent with subjects using model-based inference (Figure 1D). A closer examination of the estimated values for *G* revealed substantial variability across subjects (Figure 1E). Importantly, this parameter correlated with a purely behavioural (model agnostic) index of model-based choice (i.e. the choice consistency between generalisation trials and preceding learning trials; *r* = 0.59, *p* = 0.001; Figure 1F) but was not predicted by a measure of task transition knowledge (*r* = −0.06, *p* = 0.75; Figure 1G), indicating it reflected use of a model-based strategy rather than a readout of accuracy in deployment of task knowledge (see supplementary results).

### Model-based planning is associated with replay of task states

We first asked whether, at decision time, there was evidence for neural reactivation of task terminal goal states linked to avoidance decisions. We employed a temporal generalisation approach, as used previously^10^, to train a classifier to discriminate between the terminal states of either arm. To ensure these trained classifiers were unbiased by the state’s position or value in the task itself, we used a classifier trained on a pre-task functional localiser wherein subjects repeatedly viewed images in a random order. These images were used subsequently in the main task, prior to learning any sequential or value relationships. Thus, these classifiers captured neural responses unique to images that represented discrete states, allowing us to differentiate between the terminal state of each arm. Critically, we could apply this classifier to task data to ask if reactivation predicted behaviour (i.e. how well the classifiers predicted the chosen path at the outcome phase) Thus, if predictions from the classifier align with an actual decision, we could infer that the perceptual representation of a selected stimulus was goal-relevant at that time.

While the functional localiser included all task stimuli, for the analysis of terminal state reactivation we trained a binary classifier on these terminal states alone. Here we trained the classifier iteratively at 10ms (one sample) intervals up to 800ms post stimulus onset. Subsequently, we applied the classifier across task periods of interest at 10ms intervals, producing a two-dimensional array of classification accuracy values representing the extent to which reactivation predicted a decision on each trial.

During planning, collapsing across both trial types, we found no evidence for reactivation of terminal states, which we would expect if reactivation supported model-based planning that depends on a consideration of non-local states. However, based on prior work^8,11,15^ we reasoned that the degree of reactivation should relate to the level of model-based inference needed to make successful decisions on generalisation trials. Here reactivation is likely when making plans based on outcomes that were not the object of direct experience in relation to these arms. This predicts greater evidence for terminal state reactivation in subjects who use this model based strategy and, as detailed in Figure 1E, subjects behaviour varied in line with the extent to which they deployed model-based inference.

To test our hypothesis of greater reactivation in those deploying a model-based strategy, we examined the correlation between the predictive accuracy of our classifier (representing evidence for reactivation of the corresponding terminal states) and the generalisation parameter derived from our computational model. The latter provides a metric of the extent to which subjects used model-based inference when generalising from learned to unlearned arms. Likewise, we reasoned a positive association between model-based inference and reactivation should be more pronounced when a model-based inference is required. Thus, we conducted the analysis separately for learning trials (where no model-based inference is required) and generalisation trials (where model-based inference is required). As expected, no significant clusters were present when we examined learning trials (Figure 2A). Critically, for generalisation trials, we found a positive association between individual generalisation and choice classification accuracy during the planning period (*p* = 0.007, Figure 2B). This reflected reactivation of a component present 520ms after stimulus presentation during the localiser task. However, the interaction between learning and generalisation trials was not significant.

**Figure 2.**
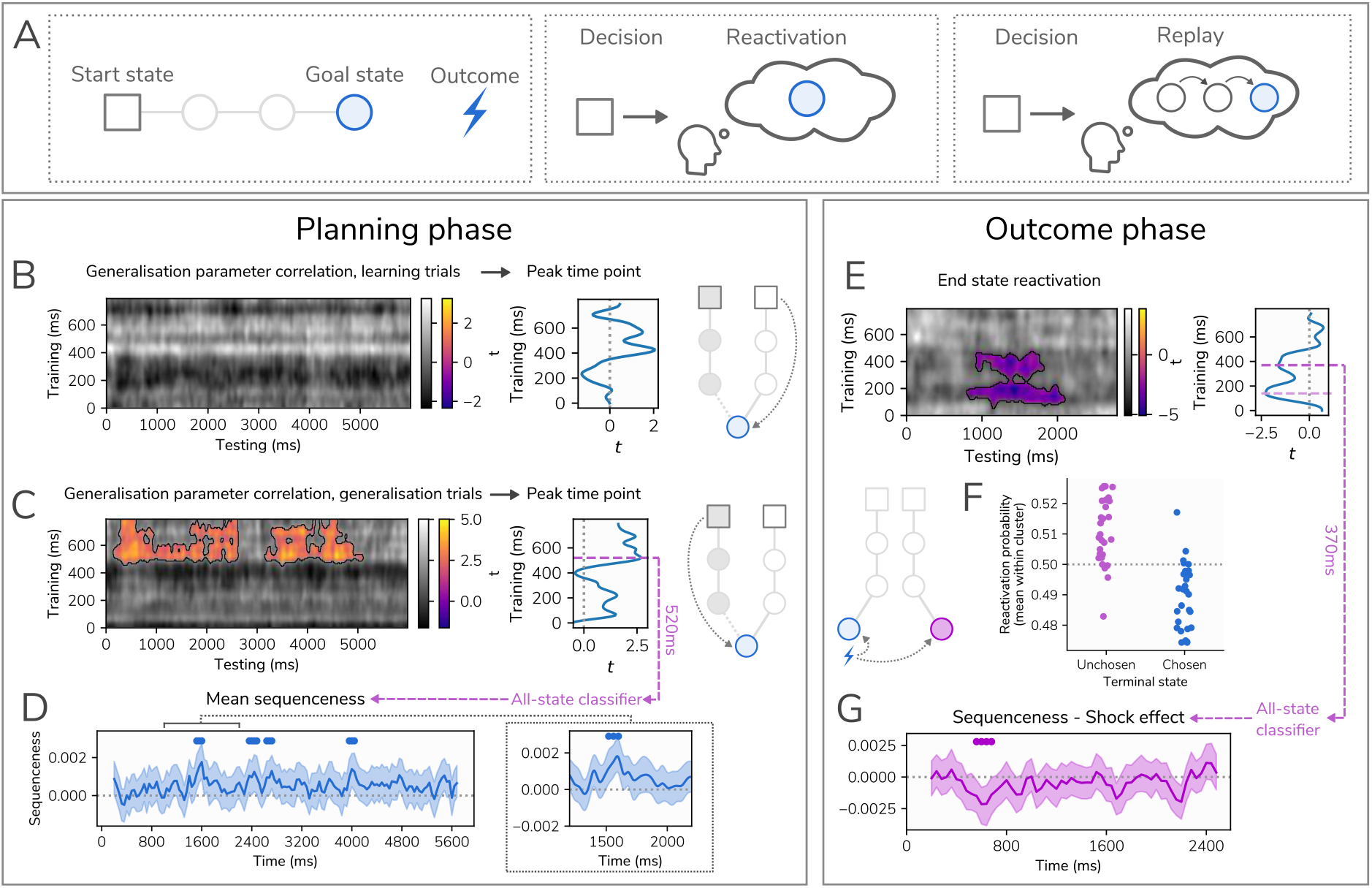
Replay and reactivation of task states. A) Illustration of approach, focusing on planning. Subjects are asked to navigate from a start state to a goal state of their choice. We assess evidence for reactivation (isolated simulation of individual states), and replay (sequential reactivation of a series of states). B) Association between classifier prediction accuracy during learning trials and the *G* parameter from our computational model, a measure of the extent to which subjects used model-based inference on generalisation trials, indicating no significant associations. C) At planning, the association between classifier prediction accuracy during generalisation trials and the *G* parameter from our computational model. Two significant clusters are seen, one from 140ms to 2860ms and one from 3150ms to 5110ms, indicating a late component of the subsequently chosen terminal state stimulus representation (peaking at 520ms) was reactivated during planning to a greater extent in subjects who used a model-based inference. D) Replay analysis during planning phase, showing mean sequenceness across all paths and trials (representing evidence for sequential replay) within the trial, based on a classifier trained at 520ms post-stimulus onset. Positive values represent forward replay, while negative values represent reverse replay. The shaded region represents the 99.9% highest posterior density interval from a Bayesian regression model, and circular marks at the top of the figure series indicate individual time points where the interval does not include zero. E) Evidence for reactivation of end states during outcome phase, showing the terminal state (either M or N) of the unchosen path is reactivated. Negative values represent classifier accuracy below chance, indicating that the classifier trained on stimulus representations is predicting the opposite state to that chosen. Two components of the stimulus representation are reactivated, one peaking at 150ms and one at 370ms. F) Mean reactivation probability across the cluster shown in (E) for end states of chosen and unchosen paths during outcome phase (learning trials only). G) Mean sequenceness based on a classifier trained on the 370ms stimulus component, where negative values indicate that shock outcomes are associated with greater reverse replay (no significant effects were seen with a classifier trained on the 520ms component). Circular markers at the top of the figure series indicate individual time points where the 99.95% highest posterior density interval does not include zero.

We asked next whether evidence for reactivation in generalisation trials embodies a sequential replay of task states, assessed using a metric termed sequenceness^14,15,28^. To address this, we used a series of 14 classifiers, each built to distinguish an individual state from all other task states (Figure S1). In contrast to the previous analysis, where we focused on preferential reactivation of terminal states alone, this classifier allowed us to test for reactivation of every task state, facilitating quantification of sequential replay of task state sequences. We trained the classifiers at 520ms post-stimulus-onset, given that this component represented the peak in reactivation in individuals who used model-based inference to a greater extent. Applying this classifier to task data provides an index of reactivation likelihood at each time point in every trial. Using Temporally Delayed Linear Modelling (TDLM)^13,14,28,28^ of lagged cross-correlations between state reactivation vectors for different states, we can determine whether reactivations occur in a sequence consistent with task structure, where positive values indicate forward replay and negative values indicate reverse replay.

While prior studies have used this approach to identify evidence for sequential replay averaging across the entire trial^15,18,28^, here we adopted a related approach used in our prior work^13,29^. The latter measures sequenceness within sliding windows that quantify fluctuations in evidence for replay within a trial. Thus, it can reveal time-varying patterns of replay, rather than relying on an assumption of a constant level of replay across the entire trial. This enables us to determine the presence of replay as well as characterize its temporal profile across the duration of a decision. In essence, instead of looking for the speed of replay (as in some prior studies), we examine the time within a trial when replay happens.

For this purpose, we tested for sequenceness, with state-to-state lags of up to 200ms, within a sliding window of 400ms across the 6 second planning period, assessing evidence for sequenceness at each time point within the trial. We used an hierarchical Bayesian latent Gaussian process regression method which accounts for a correlation between time lags (see methods). In essence, we examined modulation of sequenceness in both learning and generalisation trials by relevant task factors, assessing evidence at each time point within the trial based on the highest posterior density interval (HPDI) of the recovered Gaussian process, using a process similar to that used in our prior work on time-varying replay^30^.

On average, across trial types, all states in a path were replayed in a forward direction throughout the trial, with evidence for forward replay peaking 1600ms into the 6-second planning phase (Figure 2C). When we examined whether replay intensity differed for chosen compared to unchosen paths, we found no time points that met our criteria for significance (see methods), indicating subsequently chosen and unchosen paths were replayed to a similar extent. There was no effect of trial type, indicating that sequential reactivation was present on both learning and generalisation trials. Thus, while there was evidence of stronger reactivation of terminal states on generalisation trials, reactivation of paths leading to, and including this terminal state was no more sequential than on learning trials.

### Previously unchosen paths are reactivated during aversive learning

Theoretical work has proposed that state reactivation after feedback facilitates an outcome credit assignment that underpins value learning^31^. Here we examined this question in the aversive domain, first using the non-sequential reactivation measure associated with generalisation during planning. As at planning, this outcome phase analysis first involved training classifiers to discriminate between terminal states before using these classifiers to predict the chosen terminal state. Here we reason that If state representations are being reactivated following an outcome, this should provide sufficient information for a classifier to accurately determine which path was chosen.

We found that the accuracy of a classifier’s predictions in a cluster starting 800ms post-outcome was significantly below chance (Figure 2E, cluster *p*=.004). This indicated the classifier reliably indicated that the unchosen terminal state was reactivated following an outcome (Figure 2F). The cluster included peaks evident at 370ms and 150ms in the stimulus-locked functional localiser activity, indicating that both early and late components of the stimulus perceptual representation were reactivated, a finding reminiscent of what we had seen previously^10^.

We next asked whether post-outcome reactivation embodied a sequential component, testing whether entire paths were replayed in sequence. We again used a sliding window sequenceness analysis to identify periods in the trial where task states were reactivated in a sequence consistent with task structure. As we observed reactivation of two stimulus components (one at 150ms and one at 370ms), and prior work suggests that both early and late stimulus components represent distinct features of a stimulus^10^, we trained classifiers independently at these timepoints and evaluated evidence of sequential replay based on both, using a more conservative threshold to account for the two tests (i.e., to control for multiple comparisons). There was no evidence of sequential replay on average across shock and safety outcomes using either the 150ms or 370ms classifiers. However, average effects can conceal heterogeneity in the level of replay across trials. Decomposing an average effect on replay intensity by outcome type (shock versus safety) highlighted negative effects at several timepoints within the trial in the 370ms classifier (Figure 2H), reflecting evidence for stronger reverse replay following shock compared to safety at a cluster of timepoints beginning 600ms post-outcome.

### Medial temporal lobe theta power is linked to reactivation strength

Having established the presence of task state reactivation, we next sought to localise source regions supporting this reactivation. Given the importance of the hippocampus in reactivation and replay^15,17,32^, and the role of hippocampal theta oscillations both in memory retrieval^33,34^ and avoidance^35^, we examined whether trial-by-trial theta was associated with strength of reactivation across trials. We first performed source localisation using beamforming^36^ to provide localised estimates of theta power over the course of each trial. For the planning phase, this was limited to generalisation trials as this was where we observed evidence of reactivation. We used linear regression to predict hippocampal theta power from reactivation strength, quantified by taking probabilistic predictions from our classifiers on every trial, averaged within clusters showing evidence of reactivation (shown in Figure 2C and 2E), and performed separately for planning and outcome phases (see Methods). We performed this analysis at increasing levels of granularity, beginning by assessing predictive strength when averaging across the entire trial, then examining individual time points within the trial.

In the planning phase of generalisation trials, we found hippocampal theta power was predicted by reactivation strength in both the left (*t*(27)=3.01, *p*=.01) and right (*t*(27)=2.71, *p*=.02, Figure 3A) hippocampus. We followed up on this by asking whether activity at specific time points during the trial were especially predictive of reactivation strength. We focused on a 2000ms time window centred on the time point where we observed strongest evidence of forward replay (1600ms post trial onset), with the aim of detecting points where hippocampal activity preceded or followed replay events. Here we identified a cluster extending in time from 850-1090ms post trial onset where reactivation was significantly predictive of theta power in the right hippocampus (*p* = .02, Figure 3B). When we repeated this analysis at the whole brain level, we found no significant clusters.

**Figure 3.**
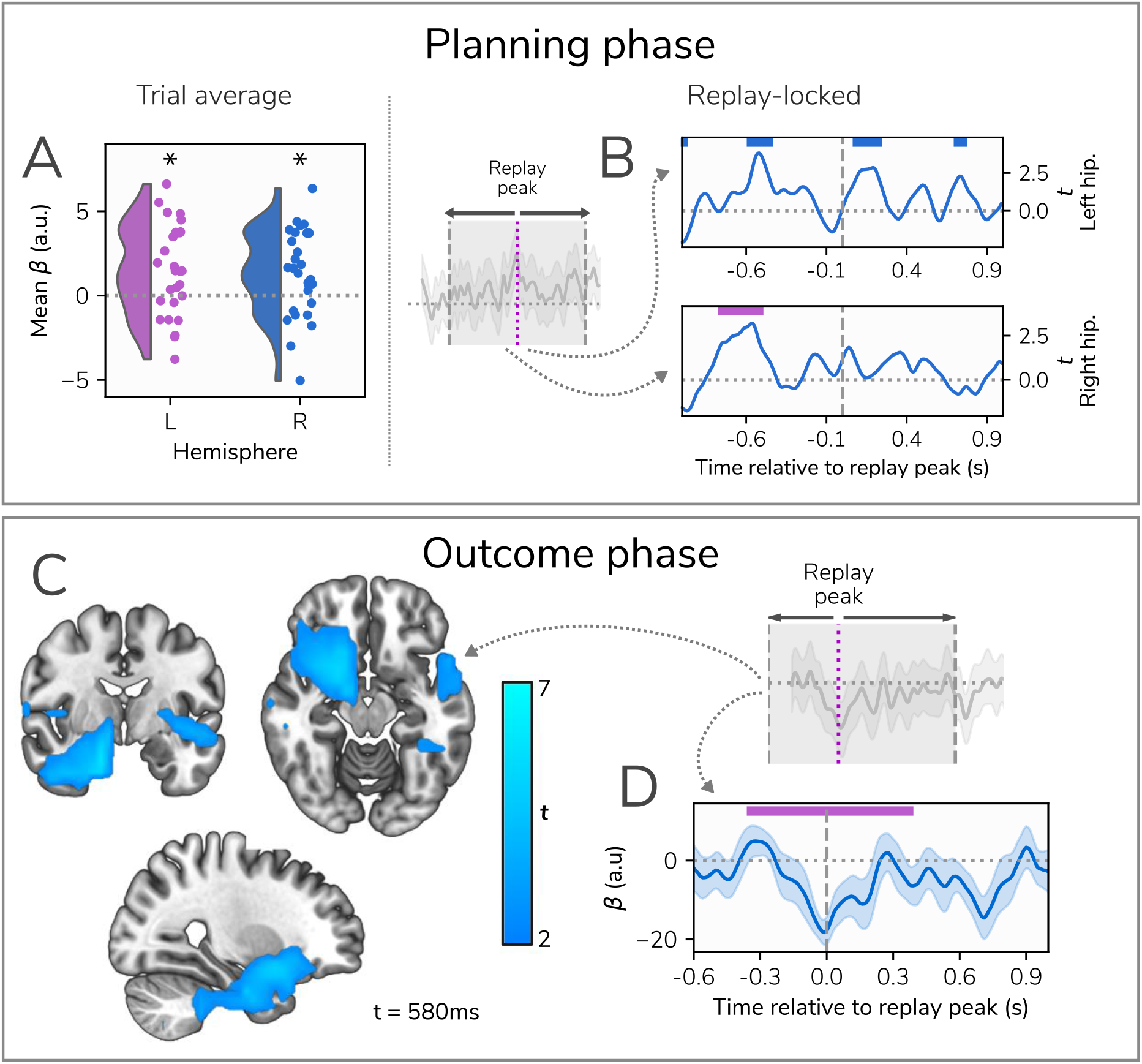
Source localisation at planning and outcome phases. A) Averaging across the entire trial, generalisation trials showed bilateral hippocampal theta power coupled to reactivation strength during the planning phase. The figure shows all subjects’ mean beta weights across the trial for the effect of reactivation on theta power, which was significantly greater than zero in both hippocampi. B) The temporal profile of associations between reactivation strength and hippocampal theta power, within a 2000ms window centred on the time point where we observed strongest evidence of forwards replay (1600ms following trial onset). The timecourses represent the effect of reactivation strength on theta power across the trial, with non-significant clusters represented by blue markers above and significant clusters represented by purple markers. C) Results of the whole brain analysis in the outcome phase, also performed within a window centred on the timepoint showing strongest evidence of replay (600ms following outcome receipt), showing a single significant cluster where theta power was negatively associated with reactivation of the chosen end state (and hence positively associated with reactivation of the unchosen end state). The image is focused on the timepoint 580ms after an outcome was displayed. D) Mean timecourse of beta weights at the peak of the significant whole-brain cluster across the trial, with the replay peak indicated by vertical dashed line and the temporal extent of the cluster shown in purple.

Next, we examined the relationship between hippocampal theta and reactivation at the outcome phase. In the hippocampus, we found no evidence for theta power predicting reactivation strength (Figure 3B), with non-significant effects across both hippocampi found when averaging across the entire trial (left *p* = 0.15, right *p* = 0.12). As with the planning phase, we next tested for significant time periods within a window centred on the time point showing strongest evidence of replay associated with shock (600ms post outcome). There were no significant time periods within the trial where hippocampal theta power was significantly associated with reactivation strength (Figure 4B). However, at the whole brain level we identified a significant cluster encompassing the amygdala, striatum, and anterior aspects of the hippocampus, and extending to encompass the ventromedial prefrontal cortex (vmPFC) where reactivation strength was negatively predictive of theta power (*p* = .01, Figure 3C). Thus, at outcome, greater theta power predicted stronger reactivation of an unchosen state that persisted in time from 260ms to 970ms post-outcome, peaking at 580ms, 20ms prior to the time point showing strongest evidence of replay (Figure 3D).

## Discussion

We show task states are replayed during aversive learning in a manner that supports model-based planning and value updating, through simulation of trajectories where direct experience of the consequence of choosing a trajectory is unavailable. These findings, consistent with rodent work, highlight a role for reactivation and replay in learning and planning during avoidance in the absence of direct experience. We speculate that replay might be a candidate neural mechanism supporting negative imagery and ruminative thought patterns.

During planning, a preferential reactivation of task goal states was seen when model-based inference was required. This depended in turn on the extent to which individual subjects actually relied upon a model-based strategy. While the difference between learning and generalisation trial types was not significant, the significant correlation of reactivation during generalisation with individual differences in model-based inference provides evidence that use of this strategy may rely on prospective reactivation. Our finding here is reminiscent of previous work in the reward domain showing that reactivation supports decisions based on model-based inference of state-value associations in subjects who demonstrate behaviour consistent with this type of strategy^8,11,15^.

Reactivation strength during planning in trials requiring model-based inference was positively associated with hippocampal theta power, with the strongest effect occurring within one second of trial onset, suggesting that reactivation is dependent on the hippocampus, as reported in rodents^37,38^. Importantly, such reactivation occurred as part of a pattern of sequential reactivation of task states in a forward direction, as found also in rodents^39^. These results implicate reactivation in avoidance planning when direct experience of outcome receipt following a specific state transition is unavailable, consistent with a simulation of outcomes and paths for actions that are yet to be taken.

After outcome receipt, reactivation of the unchosen outcome state was complemented by replay analyses where, following shock outcome, path sequences were replayed in a reverse order. Post-outcome reactivation has been proposed as a mechanism through which credit is assigned to states preceding an outcome^40^, and through which cognitive maps are updated^16,41^. We found that the reactivation of unchosen states was associated with theta power in a cluster encompassing medial temporal lobe (MTL) as well as regions such as amygdala, vmPFC and surrounding regions. While we cannot pinpoint the precise source of this reactivation using MEG alone, the regions encompassed within this cluster include those proposed to represent cognitive maps across multiple task domains^42,43^. These regions are implicated also in counterfactual thinking^44–46^ and avoidance^35,47,48^. Our results extend upon this prior work, and suggests that during aversive learning these regions support a form of counterfactual updating based on a learned model of the environment. This updating occurred rapidly after trial onset, at approximately the same time as the emergence of evidence for reverse replay, suggesting these regions initiate reactivation of relevant task states after outcome receipt.

Our results have strong parallels to findings in rodents showing preferential awake replay of paths leading to conditioned shock zones prior to avoidance^4^. Interestingly, we find that terminal states for chosen, rather than avoided, paths are reactivated. Aside from species differences, in our task subjects made deliberative avoidance plans that relied on model-based knowledge. In contrast, the prior study focused on reactive movements away from threat during free navigation^4^. Nevertheless, our results hint that similar mechanisms may support human model-based avoidance, implicating reactivation in retrieving and maintaining representations of paths towards safety and facilitating avoidance of locations associated with threat. Future work will be required to understand factors that bias reactivation towards chosen or unchosen paths in humans and other animals. In addition, while we found evidence of sequential replay during planning, its strength was only weakly related to decisions or task factors, suggesting that sequential replay may provide a readout of the task structure regardless of behaviour or current task phase.

Our work provides the first evidence of prospective and retrospective reactivation and replay during planning and learning in aversive environments in humans. Growing evidence implicates reactivation^8,10,11,49^ and replay, measured using both fMRI^17^ and MEG^14,15,18^, during learning and decision making in the reward domain. However, despite being critical for understanding common symptoms of psychopathology, the aversive domain has received little attention. One study has reported alternating reactivation of potential action outcomes during avoidance decisions^19^, suggesting that the brain simulates the outcomes of its actions during deliberation, though this latter work did not address *state* reactivation, nor model-based computation, a major focus of the current work. This is a subtle, but important, distinction as reactivation of environmental states during model-based control requires an explicit internal model of these states and the transitions between them. Our results point to similar mechanisms that are at play in both aversive and rewarding contexts, and it is also notable that replay in our task occurs online (i.e. during a task, as opposed to at rest) and within very brief time windows, as shown in recent work using non-aversive tasks^16,18^.

The approach we used allowed us to identify specific time windows wherein task states are replayed in sequence. Echoing previous results^13,29^, we found bursts of sequential reactivation at consistent time points within the trial. During planning, these peaked at 1600ms into the decision-making phase, and at 600ms following shock outcome receipt, suggesting that sequential replay is not necessarily an ongoing process but one that occurs at specific moments. This may reflect the fact that the task used here is relatively simple in terms of planning requirements, involving relatively small state and action spaces. In more complex tasks, state reactivation may persist across time to facilitate ongoing planning^14^. Another notable feature of our results is the reactivation of a relatively late component of the perceptual stimulus response (peaking at 520ms post-onset in the planning phase and 370ms in the outcome phase), which is remarkably reminiscent of our prior study involving sensory preconditioning^10^, where this reactivated component was linked to representations of stimulus category (e.g. faces or scenes) rather than immediate perceptual characteristics^10^. Although our task was not designed to dissociate different levels of stimulus representation, this timing suggests we may be detecting reactivation of a similar category-level representation in the present study.

A limitation of our task is its inability to disentangle choice and threat perception, as subjects’ decisions are guided by a desire to avoid threat, and this entails unchosen sequences are unchosen precisely because they are perceived as leading to threat. This means that we cannot determine whether reactivation is prioritised by value or by the decision making process itself. Additionally, our task to some extent confounds value and choice with experience. Aversive, unchosen paths are inevitably experienced less than safe paths, and it is possible that they are preferentially reactivated as a way of equalising experience. However, our task was designed to minimise this possibility as each path is preferable on the same number of trials, meaning that subjects choosing optimally should experience both paths approximately equally. Finally, our measure of replay relies on the subtraction of forwards from reverse replay to account for the autocorrelation in state reactivation patterns^14,15^, such that our replay measure represents a dominance of either forward or reverse replay rather than either independently.

Reactivation may be recruited to simulate state transitions and outcomes that have not, or cannot, be physically experienced. This is the case, for example, when making decisions based on transitions that cannot be directly experienced or when updating value estimates of states that were avoided. This process is relevant to understanding psychiatric symptoms linked to reactivation of negative memories, most prominently worry and rumination in anxiety and depression^7^. Notably, the content of worry and rumination is often counterfactual in nature, focusing on actions not taken^50,51^, providing at least a conceptual link to the post-outcome results reported here. Furthermore, our results implicate the medial temporal lobe, vmPFC and related areas in the initiation of state reactivation, regions implicated in depression^52–55^. By establishing a role for neural replay in aversive learning our results provide a foundation for future work that can examine how aversive neural reactivation and replay relate to specific symptoms that characterise disorders such as anxiety and depression.

## Methods

### Participants

We recruited 28 subjects from University College London subject databases who provided informed consent before beginning the study. Subjects were paid at a rate of £10 per hour, plus up to an extra £10 bonus for performance during the functional localiser task. The study was approved by the UCL research ethics committee (reference 12707/002).

### Functional localiser

The first task performed by subjects in the scanner was a functional localiser task designed to elicit neural responses specific to each stimulus used in the main task. The purpose of this was to provide data to train classifiers to decode reactivation of each image during the task itself. Each of the 14 images used in the task was shown for 0.25 seconds, being followed by two words shown on either side of the screen. One of these words described the previously shown image, and subjects had to select the correct one by pressing one of two buttons on a keypad. Following this decision, there followed a 0.25 (+/− 0.15) second intertrial interval prior to the next trial beginning. In order to boost motivation and encourage attention to the task, subjects were given a bonus payment of up to £10 that was dependent on their accuracy. Subjects completed 900 trials over 6 blocks.

### Task

Subjects completed an aversive reversal learning task in which they were instructed to avoid mild electric shocks. The task involved navigating through a map comprising 14 unique states, each state represented by a visual image (Figure 1A). The map consisted of four arms, each comprising a path of four unique states: a start state, two intermediate states, and a terminal state. Importantly, although this can be conceptualised as a spatial environment, subjects were not explicitly told of any spatial relationship between states. Each of the two end states was endowed with a probability of shock that varied according to a random walk, with the same pattern of probabilities used for every subject. Overall, each end state was safest on approximately 50% of trials, ensuring no overall bias towards one or the other being safest. Subjects were informed that these shock probabilities would change across trials, and that the two end states had independent shock probabilities.

Outside the scanner, subjects first underwent a training procedure where they learnt how to navigate successfully through states. They were first allowed to freely navigate through the two arms of the maze and “valid” transitions from the current state were indicated visually. This enabled subjects to learn which sequences of images were “valid” and which were not. After 10 trials of this free navigation, we assessed subjects’ knowledge of the task structure. In this phase, subjects were presented with the first image (shared between both sequences) and one of the two final images. They were then able to sequentially select the intermediate images that would take them from the first to the last state. Subjects progressed to the next stage if they chose correct states on 8 out of 10 trials. If this was not achieved, they returned again to the free navigation phase. This procedure repeated until the success criterion was met, ensuring that all subjects had adequate knowledge of the task structure. Crucially, this pre-training used a different set of images to the real task enabling subjects to learn the task structure without contaminating the true images with structural information.

In the main task, two arms of the map were available to select on each trial (starting with states A and D or B and C; Figure 1A). To select the arm they wished to take, subjects were asked to select a single state in the arm (which could be any state from that arm) from an array of 4 potential states, prompted by the position of that state in the sequence. Thus, although subjects were required to know the entire sequence of states for each arm, on any given trial they selected only one to indicate their choice. This served to shorten the selection time while ensuring subjects maintained an ordered representation of the state sequences, rather than representing the arms through their start states alone. For example, given a choice between arm A > E > I, M and arm D > H > L > N (see Figure 1A), and being shown the number 2 (indicating the second state in the sequence of states for each arm), subjects would choose either E or H (depending on their preference for either arm) from an array of four images including E and H.

After the functional localiser task, we calibrated a shock level for each subject. This was done prior to retraining on the task to minimise distraction. We followed a procedure used previously^25^, whereby subjects are given increasingly stronger shocks to their non-dominant hand and asked to rate their intensity on a scale ranging from 1 to 10, where 10 was the most they would be willing to tolerate. Intensity was increased until a level of 10 was reached, and this was repeated 3 times. The average of these 3 shock intensities was recorded, and we used 80% of this level as the shock intensity during the task. Following shock calibration, subjects repeated the training procedure. This was identical to the out-of-scanner pre-task training, but included the real images used in the functional localiser and task and served to teach the true transitions used for the task. Importantly, we again ensured that subjects met the pre-established criterion for structure knowledge, as our investigation focused on the use of a pre-learned map, not the learning of the map itself.

The main task is illustrated in Figure 1. Following a 6 second planning phase, subjects selected the path they wished to take using the same method as outlined for the training. Once the subject had selected a full sequence of states, the sequence was played back at 1.6 second intervals (1.2 seconds for the final state). After 1.2 seconds where the final state was displayed, the outcome was shown on an otherwise blank screen for 2.8 seconds. The type of outcome was signalled by the presence of a shock symbol (for a shock outcome) or a crossed-through shock symbol (for a no shock outcome).

The critical additional manipulation was a distinction between “learning” and “generalisation” trials. In learning trials, subjects chose between the “learning” arms of each of the two mazes. Having made a decision, they were shown the full sequence of states leading to the end state, before being shown the outcome associated with that end state. Thus, they were able to learn a shock probability associated with each of the two “learning” paths and behave accordingly using model-free learning. On generalisation trials, they made a similar choice from two “generalisation” arms and once their decision was made, the next trial began. On these generalisation trials subjects were deprived of a play-through of the selected arm states and the ensuing outcome, obviating model-free learning based upon direct experience. For learning trials, outcomes were accumulated within each of the 5 blocks, and 3 of these outcomes (either shock or safety) were chosen at random at the end of the block and administered, with a 5 second gap between outcomes involving shocks. Thus, at the end of each block, each subject received three outcomes, which were biased in favour of shock or safety depending on their choices during the task. For generalisation trials, subjects were told that the outcomes were instead accumulated throughout the task, and 10 of these would be administered at the end of the session. However, this was not actually implemented, as there was no experimental need to administer the shocks after the task. Shocks were only administered during the task itself, as the training phase only involved structure learning rather than value learning. The task had 120 trials and took approximately 35 minutes.

### MEG acquisition and processing

Scans were acquired on a CTF 275-channel axial gradiometer system (CTF Omega, VSM MedTech), sampling at 1200Hz. All pre-processing was conducted in MNE Python (http://mne-tools.github.io/)^56^. Eye-tracking data was recorded simultaneously using an Eyelink 1000 system. Data from 272 usable channels were first Maxwell filtered to remove noise, before being band passed between 0.5 and 45Hz using a finite impulse response filter. After filtering, independent component analysis (ICA) was performed on the raw data to isolate noise-related components. Components related to eye blinks were detected through both correlation with eye tracking measures and visual inspection and subsequently removed from the data, while cardiac components were detected and removed using automatic tools provided in MNE. Finally, data were downsampled to 100Hz to reduce computational burden.

### Computational modelling of behaviour

To investigate how replay relates to learning processes, we fit computational learning models to subjects’ behavioural responses. These models were Rescorla-Wagner^23^ variants, where value is updated on each trial (*t*) according to a prediction error (*δ*) weighted by a learning rate (*α*).

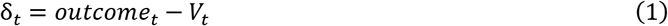

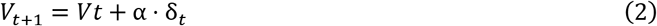

As the two end states had independent shock probabilities, the model learned the value of each separately, resulting in estimates of 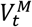 and 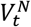 representing the two end states. Second, as the two end states had independently fluctuating shock probabilities, we adapted the update step to allow asymmetric learning, that included separate learning rates for better and worse than expected outcomes (α^+^ and α^−^ respectively).

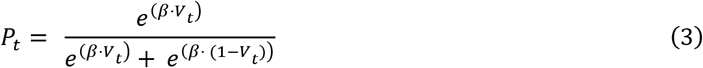

We also modified the model in order to capture choice on generalisation trials. Here, the choice was assumed to depend on the learned value of each arm’s end state (based on a generalisation of experience from learning trials). However, we allowed the degree of this influence to vary between subjects by modulating it with an additional parameter, *G*.

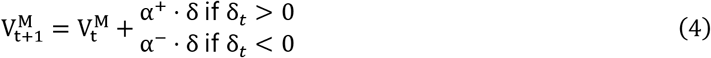

Here, *G* governs how much influence the learned end state value exerts on choice in generalisation trials. At a value of 1, these choices are fully determined by the value of the end state, and so are fully consistent (subject to decision noise) with choice probabilities on learning trials (i.e. if the subject chooses A > E > I > *M* on a learning trial with probability 90%, they are 90% likely to choose B > F > J > *M* on a subsequent generalisation trial due to the shared end state). At a *G* value of 0, choices are not influenced by learned value at all, and value is unbiased (i.e. a value of 0.5 on a scale from 0 to 1). On generalisation trials, values in the model were not updated as no outcome was presented on these trials. This model assumes that value is learned only for the terminal state in the sequence, and on learning trials, only the chosen stimulus is updated. Value is then converted to a binary choice probability using a softmax function with an estimated temperature parameter (*β*).

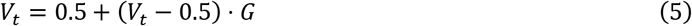

As choice probabilities did not sum to zero, they were normalised following this calculation.

All behavioural models were fit using hierarchical Bayesian methods employing variational inference implemented in PyMC3 (Salvatier et al., 2016). For further analyses involving model-derived measures, data was simulated from the model using means from each subject’s posterior distribution over each parameter value. Prediction errors were calculated by taking the difference between the expected value from the model and value of the presented outcome (1 for shock, 0 for safety).

### Temporal generalisation analyses

To perform temporal generalisation analyses, we trained binary classifiers on data from afunctional localiser to enable us to differentiate between neural representations associated with each of two given image stimuli. We built a classification pipeline using tools from MNE^56^ and Scikit-Learn^58^ that consisted of first scaling the data to have zero mean and unit variance, before performing dimensionality reduction using principal component analysis (PCA) with 50 components and then classification using L1 regularised logistic regression. The L1 regularisation hyperparameter of the logistic regression classifier was optimised using random search to produce the highest possible classification accuracy, drawing 100 potential values from a half-Cauchy distribution with *γ*=5. We trained classifiers to distinguish between start states and end states for both learning paths and generalisation paths separately, resulting in 3 classifiers (one for learning paths, one for generalisation paths, and one for the shared end states).

We applied these classifiers to the task data to determine whether they successfully predicted the path subjects chose on each trial. If the representations that classifiers were trained on were reactivated during the task in a way that relates to choice (for example reactivating state A when choosing the path that state A is a component of), then the classifiers can capitalise on this information to predict the subject’s choice. Therefore, accurate choice prediction provides evidence of reactivation of perceptual representations of task stimuli. It is important to note that the classifiers being used to predict task choices were trained on data collected prior to the actual task and therefore they are completely dissociated from any kind of value or structural associations the stimuli acquired during the actual learning task.

### State decoding for replay analyses

Data was extracted from post-stimulus onset timepoints based on those showing evidence of reactivation in the temporal generalisation analyses. These analyses were conducted using tools from Scikit-Learn^58^. Additional data was included from 50ms either side of this 150ms timepoint, such that data provided to the classifier included the period between 100ms and 200ms post stimulus onset as a form of temporal embedding. Data were subsequently subject to a dimensionality reduction procedure using temporal PCA before being transposed to an 11 (the number of timepoints) by *n* 2D matrix, where *n* represents the number of principal components. These data were then scaled to zero mean and unit variance, before being submitted to a regularised logistic regression with L1 penalty. Classification was performed using a one-vs-the rest strategy, where multiple classifiers are trained to discriminate between the stimulus of interest and all other stimuli. Sample weights were balanced to ensure classification probability was not biased by frequency of training examples. Optimal values for the number of principal components and the L1 regularisation hyperparameter of the classifier (*α)* were jointly optimised through 100 iterations of randomised search^59^ to identify the settings producing the highest classification accuracy for each subject, with each iteration of the search evaluated using three-fold cross-validation. The number of components was sampled from a uniform distribution *unif* (30, 60), while *α* was sampled from a half-Cauchy distribution with *γ*=5.

### Replay analysis

To index reactivation of task states, during planning and task rest periods, the classification pipeline trained on each subject’s functional localiser data was applied to the data to produce a probabilistic prediction of the state being activated at each time point. This involved extracting data for 50ms prior to and following the timepoint of interest, applying the PCA decomposition and scaling determined using the localiser data, multiplying the data by the trained feature weights to produce a probabilistic prediction of which state was activated at the specified timepoint. This resulted in a matrix representing the probability of each state being reactivated at each of 590 timepoints (the 600 available timepoints minus the first and last 5 timepoints, which are lost due to the temporal embedding procedure).

To detect sequential reactivation of state representations in this data, we used a cross correlation method^13,14,29^. While this method does not allow us to investigate forward and reverse replay independently due to the high autocorrelation present in the signal (instead w use the difference between these metrics to obviate this autocorrelation), the cross-correlation method is more amenable to windowed analyses than general linear modelling approaches^28^. Here, we took the product of the state X time reactivation matrix and the task transition matrix, where the time course of each state’s reactivation probabilities was correlated with the time courses for all other states, lagged by up to 20 timepoints (representing 200ms) in steps of one. This produced a matrix of correlation coefficients representing the effect of past reactivations (across 20 lags) on current state reactivation, and hence the correlation between past activation and present activation, within the transition matrix of the task. This allowed us to isolate replay of transitions that were consistent with the transitions in the task (representing forward replay) or reverse transitions (representing backward replay). These were then subtracted to produce a difference measure representing the balance of forward versus reverse replay. We assessed the transition matrices representing each arm separately, resulting in vectors of evidence for replay (termed sequenceness) across each time lag (up to 200ms) for each arm. In order to assess how sequenceness fluctuates within-trial, we performed this procedure within 400ms sliding windows, as in prior work^13,29^. This provided an index of sequenceness at each time point within the trial.

### Statistical analysis of sequenceness data

Our sequenceness detection procedure provided a measure of sequential replay at each time point, within each trial, for each subject. Our aim was to identify periods during the trial where we observe consistent evidence for sequential replay across subjects and examine whether replay is modulated by task factors (such as path type and choice). To achieve this, we used hierarchical Bayesian latent Gaussian process regression (HLGPR). This builds upon prior work using hierarchical Gaussian processes (GPs) to directly model observed timeseries, both in MEG work^13^ and other fields^60,61^, and provides an intuitive approach to modelling timeseries data while accounting for its covariance structure. This is particularly appealing for the analysis performed here, as directly accounting for covariance allows us to circumvent multiple comparison corrections across time points, which presents a problem when identifying significant effects if we expect these effects to be brief in their duration.

Here, we extend on this prior implementation to allow more flexible analysis of time-varying modulation of the observed time course. Rather than directly representing the observed timeseries as a GP, the HLGPR approach instead represents the observed timeseries as a linear combination of input features and time-varying regression weights, which are modelled as latent GPs. As these latent GPs represent a distribution over functions, we can use Bayesian modelling to approximate the posterior distribution over potential functions linking task features to replay over time.

We define this as a Bayesian linear model, where the observed data at each time point *t* is modelled as a normal distribution with mean defined as the combination of regressors and regression weights at each time point, and a single estimated variance across all time points σ.

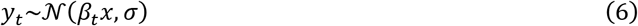

The mean of this distribution is calculated using a standard linear regression model for regressors 1,…,*n*:

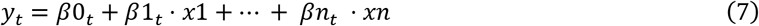

Here, each β_*t*_ is one point on a function described by a latent GP, resulting in regression weights being represented as GPs of the same length as *y*, one per regressor plus an intercept. The regressors themselves are assumed to be constant for the duration of the trial (representing a variable such as trial number, or choice on the trial). We then extend this model to allow both random slopes and random intercepts, whereby observed data for each timepoint *t* within each subject *i* = 1,…, *n subjects* is represented by the following regression model:

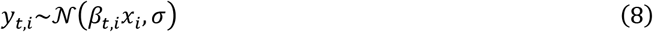

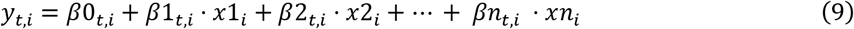

This results in a separate function per each subject for each regressor, allowing the time course of each regressor’s effect to vary across subjects. However, rather than treating each subject independently, we represent the data using a hierarchical model where we account for covariance between subjects. This approach allows us to take advantage of the benefits of hierarchical models in terms of aiding parameter estimation when using data that is limited in quantity and quality, as is typical of neuroimaging data. To achieve this, we turn to prior work on hierarchical GPs^60,61^, and use a hierarchical mean and covariance structure to define the GPs used to represent regression coefficients. For each regressor, we first define a constant-mean group-level GP representing the group mean time-varying regression weights. For consistency with prior work, we use x to represent each time point.

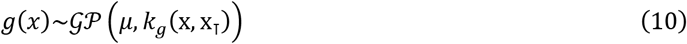

The covariance function *k*_*g*_(x, x_⊺_) is flexible, depending on the expected covariance structure of the data. Here we used the squared exponential kernel, which is parameterised by a length scale parameter *ℓ*_*group*_, for which we use a gamma prior (γ(3, 1)). We also estimate the variance of the kernel using an additional free parameter *σ*_*group*_, which is given a gamma prior (γ(3, 5)).

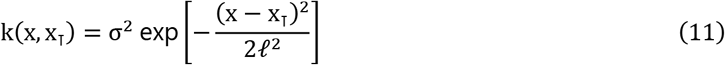

We place a normal prior 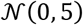 on the mean at the group level (μ), allowing the regressor to have a constant effect across all time points within the trial. We assume that each subject’s function for each time-varying regression weight is offset from this by a degree determined by a second, subject-level offset function, modelled as another GP. Thus, each subject’s regression weights are represented as the summation of the group-level function and their own subject-level function. Importantly, this subject-level function represents the degree to which their own function is *offset* from the group function, rather than representing the subject’s own function itself. This is defined as a GP with mean *g*(*x*), representing the group-level effect, and a subject-level covariance function, for which we again use a squared exponential kernel with an estimated subject level variance parameter *σ*_*subject*_ and length scale parameter *ℓ*_*subject*_.

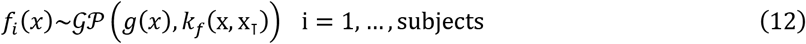

As GPs are nonparametric, the only parameters we estimate are the length-scale parameters of the covariance functions. For conceptual simplicity and computational ease, we assume the same length scale parameter across subject and group level GPs but allow this to vary across regressors.

This approach provides us with group-level GPs that represent the effect of a given regressor on sequenceness at each time point in the trial. This approach allows a far greater amount of input data to be considered than modelling timeseries as observed GPs. For example, here we are able to use sequenceness timeseries for each trial, rather than averaging across trials as in prior work ^13^. This both provides a far richer source of input data and allows us to explore the effects of continuous regressors. Using a hierarchical formulation also allows us to take advantage of the properties of hierarchical models that permit more accurate parameter estimation, while taking a Bayesian approach allows us to approximate the full posterior over parameters, and hence the posterior over functions. As a result, we can extract both the estimated mean effect of a given regressor and the 99.9% highest posterior density intervals (HPDIs) at each time point as a conservative range of likely values for the true function, allowing us to judge whether we can be confident that it has a non-zero effect at any given time point without the need for multiple comparison correction. For analyses in the outcome phase, we used the 99.95% HPDI to account for the two time points used to train classifiers. Validation analyses are presented in Supplementary Material, demonstrating that this method successfully controls the false positive rate. However, to provide additional protection against false positives, we only interpret time points that were part of a cluster of three or more temporally contiguous time points with HPDIs that did not include zero.

For the planning phase, we included regressors representing trial type (learning or generalisation), path type (learning or generalisation); whether or not the path was chosen; the interaction between trial type and path type; and trial number. For the outcome phase, we included regressors representing whether or not the path was chosen, the path type, the outcome (whether or not a shock was received), the absolute prediction error, the interaction between outcome and absolute prediction error, and the trial number.

These models were constructed and fit in Python using PyMC3^57^, and we approximated posterior distributions of model parameters using variational inference. To ease the computation burden of this analysis we decimated the data by a factor of 4, reducing our effective sample rate from 100hz to 25hz.

### Source localisation analysis

Source localisation analyses were performed using the MNE-Python implementation of the unit noise gain linear constrained minimum variance (LCMV) beamformer method. Data from task periods of interest (planning and outcome phases) were first filtered between 4 and 8hz to retain only activity in the theta band using a finite impulse response filter. The head model used to generate the source space was the standard Freesurfer subject (“fsaverage”) with a 5mm grid, and the forward solution for each subject was calculated using this head model scaled according to subject-specific transform parameters, determined manually by aligning subjects’ head position coils with those of the standard subject. The covariance of the data was estimated based on the full duration of the trial, while noise covariance was estimated based on sensor data from 500ms pre-outcome to outcome onset (this was used for both planning and outcome phases). Covariance matrices were regularised by a factor of 0.05. This procedure provided source activity estimates for each epoch. As we were interested in theta power, we finally applied a Hilbert transform to the source estimates (already restricted to the theta band) to provide an index of instantaneous power in the theta band. We chose to focus on the bilateral hippocampus, and therefore extracted mean power timeseries from these regions (defined according to the Freesurfer atlas) for our primary analyses. However, we also performed exploratory whole-brain voxel-level analysis to identify effects outside our regions of interest.

### Source-level prediction of reactivation strength

To identify associations between theta power at the source level and reactivation strength, we used a general linear model predicting theta power from reactivation strength across trials, in addition to an intercept term, at each timepoint within the trial. Reactivation strength was calculated using probabilistic predictions from our classifiers trained to distinguish between states of interest in our temporal generalisation analyses. These probabilistic predictions can be interpreted as providing an index of the strength of reactivation on a given trial (for example, if the classifier predicts that state A is reactivated with 90% probability, we take this as strong evidence for reactivation of state A). As we train classifiers at multiple timepoints in the functional localiser and apply them across multiple timepoints in the task, we generate a matrix of predictions for each trial. To obtain a single value per trial indicating strength of reactivation, we extract the mean predicted reactivation probability within a region of interest based on clusters showing significant effects in the temporal generalisation analysis. The linear model was then constructed as follows:

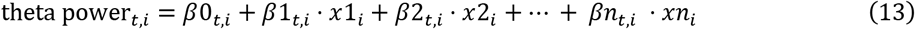

Where *t* represents the time point in the trial, *i* represents the trial number, and *x* represents a regressor. For the outcome phase, we additionally included regressors representing the outcome of the trial (shock or safety), the absolute prediction error (derived from the computational model), and the interaction between outcome and absolute prediction error (representing a signed prediction error). All regressors were mean-centred prior to being entered into the model. This resulted in a timeseries of β coefficients for each regressor at each time point in the trial. This procedure was performed for each subject using mean extracted theta power from both hippocampi, and at the voxel level in whole-brain analyses. Before aggregating results across subjects, we first standardised the *β* coefficients by dividing them by their variance to ensure they remained on a consistent scale.

We assessed significance in three steps. First, we examined the average association between theta power and reactivation across the trial by taking the mean of the *β* timeseries for each subject. These mean *β* values were then compared against zero using a one-sample t-test, with *p* values subject to Bonferroni correction for two comparisons (left and right hippocampus). Second, we sought to identify periods in the trial where theta power was most strongly associated with reactivation. Here we used one-sample permutation tests on beta timeseries against zero with sign-flipping to identify significant clusters in time (based on cluster mass). Third, we examined whole-brain results using four-dimensional cluster-based permutation tests to identify significant clusters across space and time. Beta maps were first thresholded at *p* < .01 voxelwise to identify clusters, which were then tested for significance using permutation methods based on their mass across space and time.

## Acknowledgements

T.W. is supported by a Wellcome Trust Sir Henry Wellcome Fellowship (206460/17/Z). R.J.D. holds a Wellcome Trust Investigator award (098362/Z/12/Z). Y.L. is supported by the Max Planck Society (MPS). The Max Planck UCL Centre is a joint initiative supported by UCL and the Max Planck Society. The Wellcome Centre for Human Neuroimaging is supported by core funding from the Wellcome Trust (203147/Z/16/Z). We thank Helen Schmidt and Daniel Bates for assistance with data collection.

