## Supplementary material for "Model-based aversive learning in humans is supported by preferential task state reactivation"

[*]These authors contributed equally

### Supplementary results

#### Model fitting

To verify that the *G* parameter in our model captured the kind of model-based choice we intended, we correlated it with an approximate behavioural index of generalisation, namely choice consistency between learning and generalisation trials. If subjects are indeed generalising based on learned value, their choices on generalisation trials should approximate those on learning trials immediately preceding these trials. In line with this, our generalisation parameter strongly correlated with the proportion of trials where generalisation choices were consistent with preceding non-generalisation choices for each subject ($r$ = 0.59, $p$ = 0.001, Figure 1F), indicating this parameter provides a valid index of generalisation. Finally, it is possible that subjects may use model-based inference due to poor general task knowledge, rather than absence of a model-based generalisation *per se*. To ascertain this was not the case, we examined the relationship between generalisation parameter values and the number of mistakes subjects made on generalisation trials (i.e. not selecting any option or making an incorrect state selection). If subjects do not generalise due to poor task knowledge, we would expect a negative correlation between these variables. These two variables were not correlated (r = -.06, $p$ = 0.75, Figure 1F), indicating that low generalisation parameter values are not simply a reflection of poor task knowledge or execution.

#### Classifier accuracy

To determine the accuracy of the 14-way classifiers used for determining state reactivation in the sequenceness analyses, we compare the accuracy of predictions in the localiser data, using 5-fold cross-validation. Confusion matrices, collapsed across subjects, are shown in Figure S1 for each time-point used for training. Classification accuracy was above chance (1/14 = 7.14%) for every stimulus.

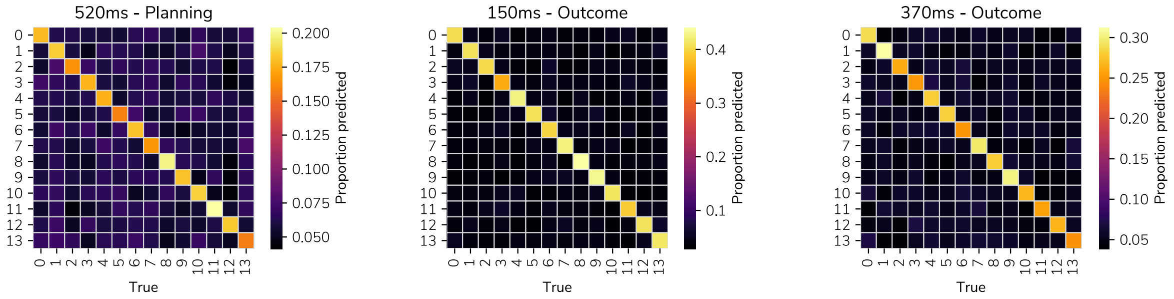


*Figure S1. Confusion matrices representing classification accuracy across different training timepoints. Values show accuracy collapsed across subjects.*

#### Validation of hierarchical latent Gaussian process regression

To validate the statistical approach used for testing time-varying sequenceness, we evaluated its performance on simulated data. The method itself is described in full in the Methods section. We generate data according to a regression model where the regressors are represented by time-varying functions. Thus, the sequenceness value on each trial is represented by the value of that trial on a given variable (this may be trial number, for example), multiplied by its regression coefficient. As we expect these regression coefficients to vary over the course of the trial (for example the effect of trial number may be greatest at the start of the trial) with a degree of autocorrelation, representing the regressors as functions of time provides a simple model of the data-generating process. We used 100 simulated trials, and 100 timepoints.

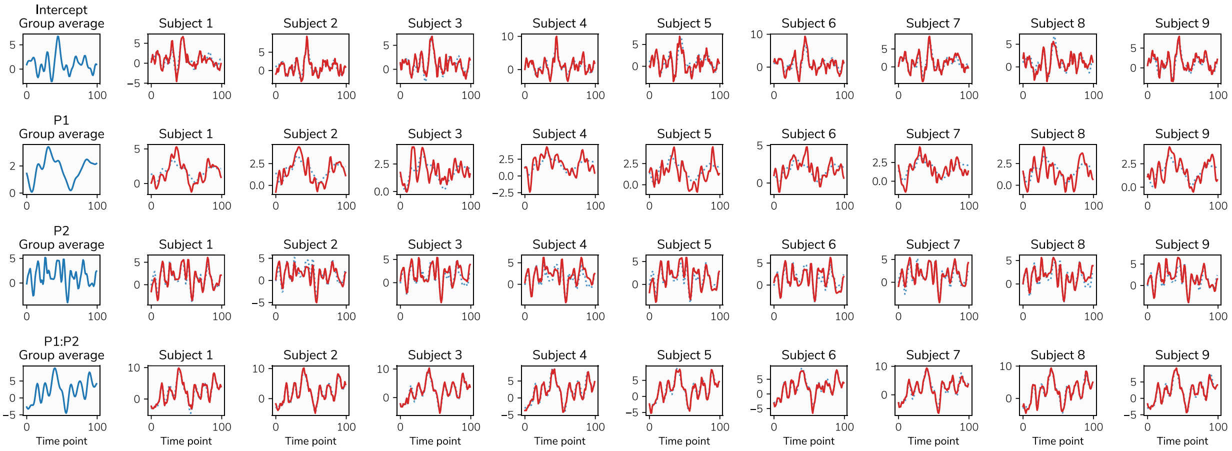


*Figure S2. Functions used to generate simulated data. The blue lines on the left of the figure are the group-level functions. The red lines on the right side are the subject-level functions, with the group-level function shown in the dotted blue line.*

For the purpose of validation, we simulate data from four known functions determined by Gaussian processes with pre-determined covariance functions, with the first representing an intercept, two representing separate regressors, and the final representing the interaction between the previous two regressors. We structure the data in a hierarchical manner, with these functions representing the group-level process. The covariance function of each Gaussian process has its own length scale parameter (values of 3, 5, 2, 4), and its own variance (2, 0.8, 2, 3). The mean function of these Gaussian processes is set to a constant (with values of 1.5, 2, 0.3, 0.7), representing an effect that does not depend on time.

We then determine 20 subject-level functions (assuming 20 simulated subjects) for each regressor, which are represented by the sum of the group level function and a separate, subject level function. Each subject-level function is drawn from another Gaussian process, again with separate length scale (3, 5, 2, 4) and variance (1, 0.8, 0.9, 0.8) parameters for the covariance function. Thus, each subject's function is offset from the group-level function by a subject-specific function. The length scale parameters represent the degree of correlation between adjacent timepoints, while the variance parameters have the effect of representing the range of the function. A greater subject-level variance relative to the group-level variance has the effect of increasing the subject-level variability and reducing the influence of the group-level function. Examples of these functions are shown in Figure S2. Finally, to represent error, we add noise determined by a Gaussian distribution with zero mean and a standard deviation of 20. We also repeated this process using simulated data with no effect at the group level; that is, subjects had their own effects but these were offset from a group effect that was zero across all time points.

Fitting the models of the form used for the sequenceness analyses to this simulated dataset allowed us to recover the true group-level functions used to generate the data (for the purposes of these analyses, we are interested only in the group-level effect). As in the analyses reported in the main results section, we used the 99.9% highest posterior density intervals (HPDI) of the posterior distributions over these functions to obtain a conservative metric through which to determine whether the recovered interval includes the true function. Calculating the number of timepoints where the HPDI includes the true function value provides a measure of the false positive rate. The results of this test are shown in Figure S3.

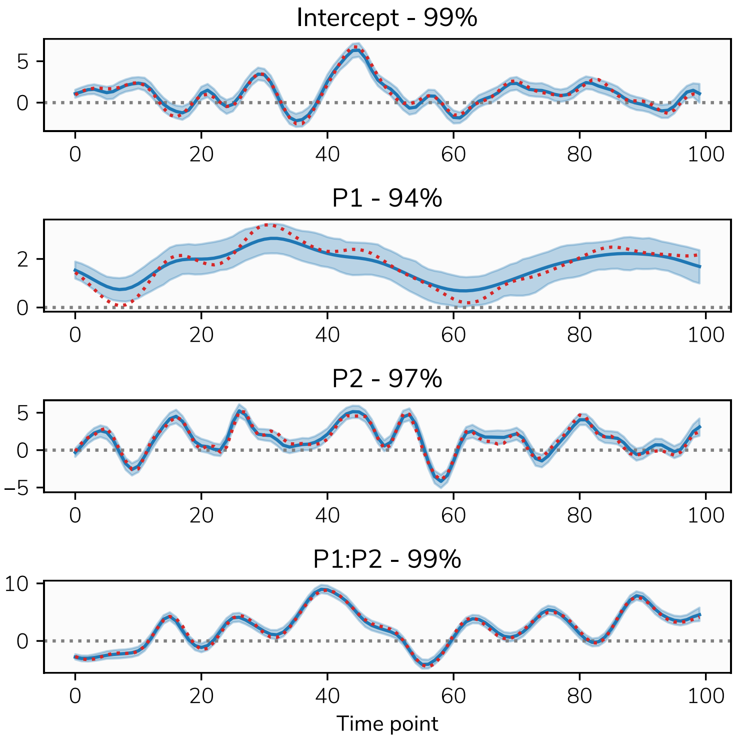


*Figure S3. Means and 99.9% HDPIs of the recovered group-level function, shown in blue. The red dotted line represents the true group-level function.*

As shown in Figure S2, the true function was included in the HPDI in 97.25% of time points across the four regressors, indicating that while the HDPI is not perfectly calibrated with respect to the false positive rate, using the conservative 99.9% HPDI provides acceptable control of the false positive rate. As shown in Figure S4, repeating this with a null effect at the group level (i.e. an effect of zero across all time points), the 99.9% HPDI includes zero at 100% of time points.

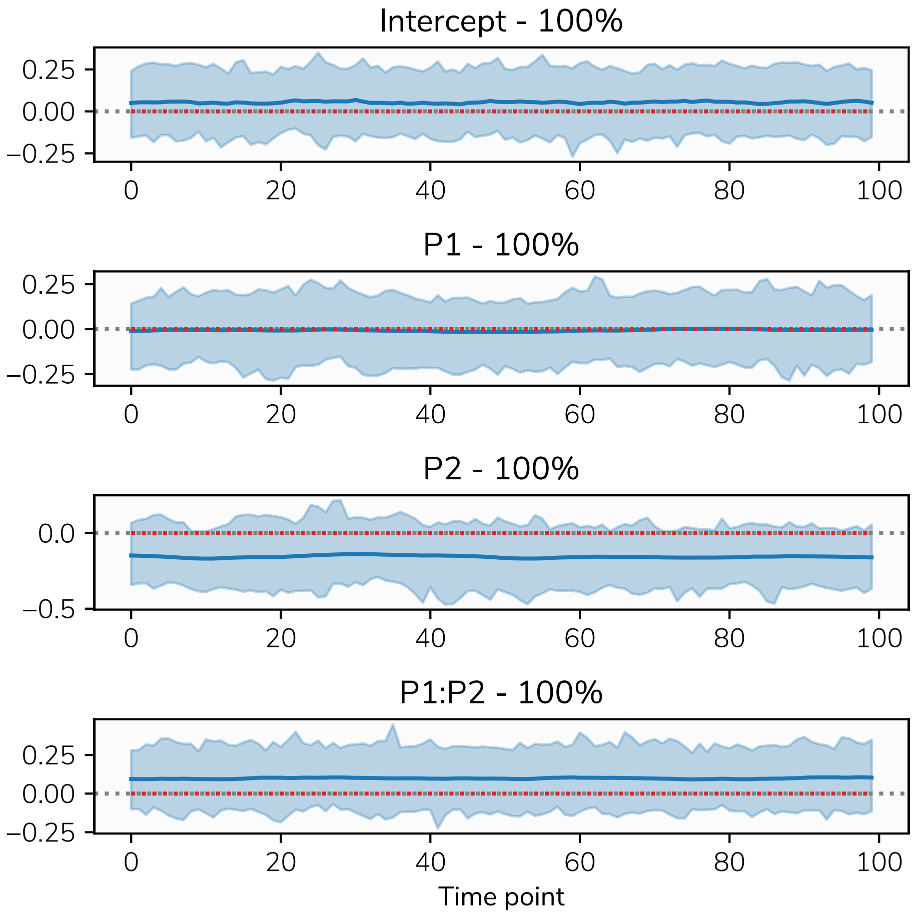


*Figure S4. Means and 99.9% HDPIs of the recovered group-level function (showing no effect across the trial), shown in blue. The red dotted line represents the true group-level function (zero across the entire trial).*
